# Biologically relevant integration of transcriptomics profiles from cancer cell lines, patient-derived xenografts and clinical tumors using deep learning

**DOI:** 10.1101/2022.09.07.506964

**Authors:** Slavica Dimitrieva, Rens Janssens, Gang Li, Artur Szalata, Raja Gopal, Chintan Parmar, Audrey Kauffmann, Eric Y. Durand

## Abstract

Cell lines and patient-derived xenografts are essential to cancer research, however, the results derived from such models often lack clinical translatability, as these models do not fully recapitulate the complex cancer biology. It is critically important to better understand the systematic differences between cell lines, xenografts and clinical tumors, and to be able to identify pre-clinical models that sufficiently resemble the biological characteristics of clinical tumors across different cancers. On another side, direct comparison of transcriptional profiles from pre-clinical models and clinical tumors is infeasible due to the mixture of technical artifacts and inherent biological signals.

To address these challenges, we developed MOBER, **<u>M</u>**ulti-**<u>O</u>**rigin **<u>B</u>**atch **<u>E</u>**ffect **<u>R</u>**emover method, to simultaneously extract biologically meaningful embeddings and remove batch effects from transcriptomic datasets of different origin. MOBER consists of two neural networks: conditional variational autoencoder and source discriminator neural network that is trained in adversarial fashion. We applied MOBER on transcriptional profiles from 932 cancer cell lines, 434 patient-derived tumor xenografts and 11’159 clinical tumors and identified pre-clinical models with greatest transcriptional fidelity to clinical tumors, and models that are transcriptionally unrepresentative of their respective clinical tumors. MOBER can conserve the biological signals from the original datasets, while generating embeddings that do not encode confounder information. In addition, it allows for transformation of transcriptional profiles of pre-clinical models to resemble the ones of clinical tumors, and therefore can be used to improve the clinical translation of insights gained from pre-clinical models. As a batch effect removal method, MOBER can be applied widely to transcriptomics datasets of different origin, allowing for integration of multiple datasets simultaneously.

## Introduction

Cancer cell lines and patient-derived tumor xenograft models continue to play a critical role in pre-clinical cancer research and drug discovery^1–3^. Thousands of cancer models have been established and propagated *in vitro* and *in vivo* in different laboratories, where they have been extensively used in pre-clinical studies to study the biology of cancer^4,5^, to explore the vulnerabilities of cancer cells^6,7^, to identify new biomarkers^8^ and to test the efficacy of anticancer compounds^2,9^. Enormous knowledge in cancer biology has been derived from the various experiments conducted on cancer models, still, many of the significant findings from pre-clinical cancer research are not reproducible in clinical trials^10,11^ and oncology drugs have the highest failure rate compared to compounds used in other disease areas^12^. One of the major reasons to this lack of translatability is that the cancer models are not perfect, and because of their propagation and differences in growing conditions, they have altered over time, and it is not known how well they represent the biology of the tumors from which they were derived. In addition, many cancer models lack accurate clinical annotations and histopathological classification, that are crucial for their utility in cancer research^13^. For greater clinical translatability, identification of models that sufficiently resemble the biological characteristics and drug responses of patient tumors is of critical importance.

Large collections of molecular data from patient tumours and cancer models have been generated across different cancer types. The *Broad-Novartis Cancer Cell Line Encyclopedia* (CCLE)^2^ contains molecular profiles of around thousand cancer cell lines, which are extensively used as pre-clinical models for various tumour types in drug discovery studies. In addition, gene expression profiles of >400 patient-derived tumor xenograft (PTX) models are available via the *Novartis Institutes for Biomedical Research Patient-derived Tumor Xenograft Encyclopedia*^9^. Comprehensive molecular characterization of primary and metastatic tumors along with clinical data from >11,000 patients are available from the *The Cancer Genome Atlas* (TCGA)^14^, MET500^15^ and *Count Me In* (CMI)^16^ projects. These efforts provide a powerful opportunity to unravel the systematic differences between cancer cell lines, xenograft models and patient tumors, and to identify the cancer models that sufficiently recapitulate the biology of patient tumors without relying on clinical annotations.

Gene expression profiling accurately reproduces histopathological classification of tumors and is a useful technique for resolving tumor subtypes^17–20^. However, large-scale integration of molecular data from cancer cell lines, xenograft models and patient tumors is challenging due to the mixture of intrinsic biological signals and technical artifacts. One key challenge is that gene expression measurements from bulk patient biopsy samples are confounded by the presence of human stromal and immune cell populations, that are not present in cancer models. In addition, large public datasets can be confounded by hidden technical variables, even when they come from the same source type (e.g. RNAseq from different patient cohorts). Existing approaches for removing batch effects do not account for other systematic differences between cancer models or patient tumors, or assume that the cell line and tumor datasets have the same subtype composition^21,22^. Previous studies analyzing the differences between cell lines and patient tumors based on transcriptomics profiles have primarily focused on selected cancer types^23–25^. Existing global analysis with the Celligner method^19^, which leverages a computational approach developed for batch correction of single-cell RNASeq data, has compared cancer cell lines and patient tumors. However, this method is limited to aligning only two datasets simultaneously and the Celligner alignment does not consider patient-derived tumor xenografts.

Here, we introduce a novel deep learning-based method, MOBER (**<u>M</u>**ulti-**<u>O</u>**rigin **<u>B</u>**atch **<u>E</u>**ffect **<u>R</u>**emover), that performs biologically relevant integration of pan-cancer gene expression profiles from cancer cell lines, patient-derived tumor xenografts and patient tumors simultaneously. MOBER can be used to guide the selection of cell lines and patient-derived xenografts, and identify models that more closely resemble patient tumors. We applied it to integrate transcriptomics data from 932 cancer cell lines, 434 patient-derived tumor xenografts and 11’159 patient tumors from TCGA, MET500 and CMI, without relying on cancer type labels. We developed a web application that provides a valuable resource to help researchers select pre-clinical models with greatest transcriptional fidelity to clinical tumors. MOBER can be broadly applied as a batch effect removal tool for any transcriptomics datasets and we made the method available as an open-source Python package.

## Results

### The MOBER method

MOBER is an adversarial conditional variational autoencoder that generates biologically informative gene expression embeddings robust to confounders. MOBER consists of two neural networks (see Methods for more details). The first is a conditional variational autoencoder (VAE)^26,27^ that is optimized to generate embeddings that can reconstruct the original input. The second is an adversarial neural network (aNN)^28^ that takes the embedding generated by the VAE as an input and tries to predict the origin of the input data. The VAE consists of an encoder that takes as an input a gene expression profile of a sample and encodes it as a distribution into a low dimensional latent space; and a decoder that takes an embedding sampled from that distribution and the origin sample labels to reconstructs the gene expression profile from it. The goal is to learn an embedding space that encodes as much information as possible on the input samples, while not encoding any information on the origin of the sample. To achieve this, we train the VAE and aNN simultaneously. The VAE tries to successfully reconstruct the data while also preventing the aNN from accurately predicting the data source. This way, during training, the VAE and the aNN will converge and reach an equilibrium, such that the VAE will generate an embedding space that is optimally successful at input reconstruction and the aNN will only randomly predict the origin of the input data from this embedding. In other words, the VAE will converge to generating an embedding that contains no information about the origin of the input data, while the aNN will converge to a random prediction performance. During the MOBER training process, we use one-hot representation of the origin of the input samples (either TCGA, CCLE, PTX, MET500 or CMI) and provide that information to the decoder. This enables the VAE to decode the data embeddings conditionally, and reconstruct the expression data as if the sample was coming from another source type. To project a transcriptomics profile from one origin (e.g. pre-clinical) into another (e.g. clinical), after the model is trained, we can pass that transcriptomics profile through the VAE and simply change the one-hot vector informing on the sample origin to the desired one (e.g. CCLE to TCGA to decode cell lines as if they were TCGA tumors).

### Global pan cancer alignment of transcriptional profiles from cancer cell lines, patient-derived tumor xenografts and patient tumors

We analyzed the transcriptomics profiles from 932 CCLE cell lines, 434 patient-derived tumor xenografts, 10’550 patient tumors from TCGA, 406 metastatic tumors from MET500^29^ and 203 breast tumors from *Count Me In* (CMI)^16^. Integrating these datasets by performing dimensionality reduction with 2D Uniform Manifold Approximation and Projection (UMAP), reveals a clear separation of samples based on their origin (Fig. 2a). As expected, there is a global separation between cell lines, xenografts and patient tumors. In addition, there are still strong batch effects between CMI, TCGA and MET500 datasets, despite these samples all being derived from patient biopsies. This separation was not addressed by applying *ComBat*^21,22^, a widely used batch correction method (Suppl. Fig. 1).

**Fig. 1.**
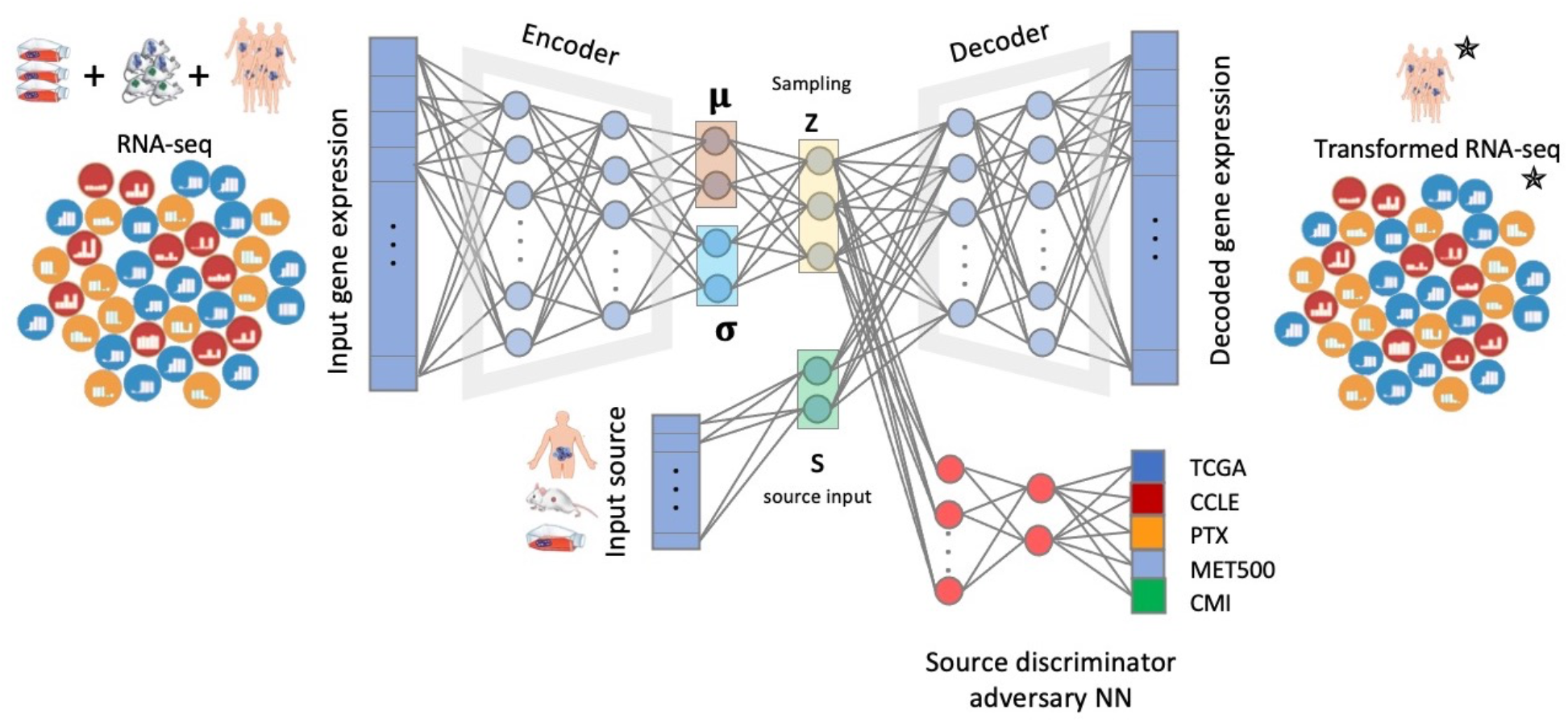
MOBER architecture. MOBER consists of two neural networks: conditional variational autoencoder and source discriminator neural network that is trained in adversarial fashion. The encoder takes as an input gene expression profile and encodes it into a latent space, the decoder takes a sampling from the latent space and reconstructs the gene expression profile from it. The source discriminator adversary neural network takes the sampling from the latent space and tries to predict the origin of the input data.

**Fig. 2.**
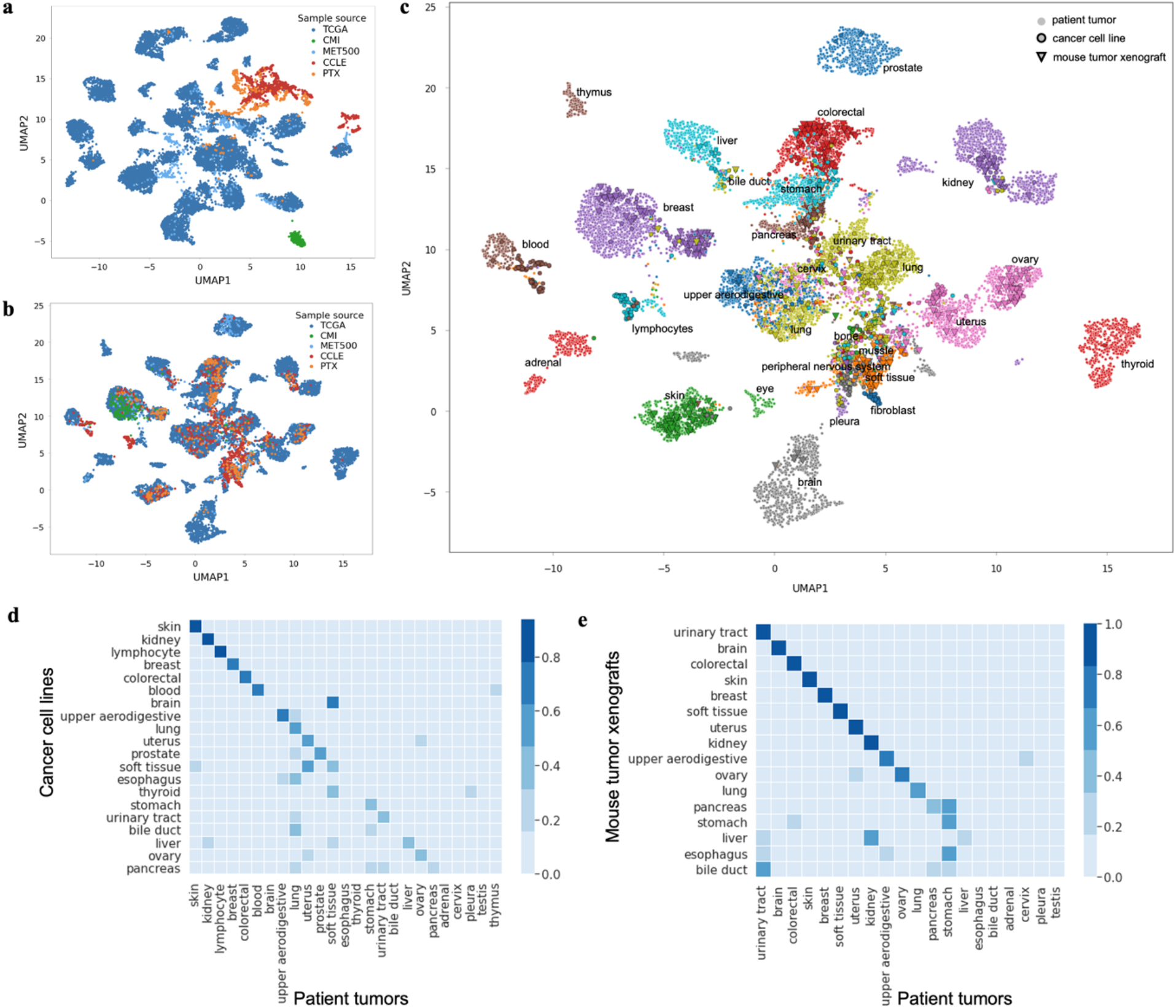
Global pan cancer alignment of pre-clinical and clinical transcriptomes. a) Integration of transcriptional profiles from models and patient tumors by performing UMAP dimensionality reduction, each dot is a sample. b) Integration of transcriptional profiles from models and patient tumors using MOBER, the color corresponds to the sample origin. c) MOBER alignment, where each tumor sample is colored based on cancer indication. d) The proportion of cancer cell lines that are classified as each tumor type using MOBER-aligned data. e) The proportion of PTXs that are classified as each tumor type using MOBER-aligned data. The x-axis on d) and e) shows the TCGA tumor types, the y-axis shows the CCLE annotation label (d) and PTX annotation label (e), accordingly.

Aligning the CCLE, PTX, TCGA, MET500 and CMI datasets with MOBER resulted in a well-integrated dataset of transcriptional profiles from cell lines, xenografts and patient tumors that have been corrected for multiple sources of systematic dataset-specific differences. Indeed, the UMAP plots using the MOBER-aligned data (Fig 2b,c) reveal a map of cancer transcriptional profiles where cell lines, xenografts and patient tumor samples are largely intermixed, while the biological differences across known cancer types are still preserved (Fig. 2c).

As illustrated in Fig. 2b, MOBER removes the systematic differences between patient tumors and cancer models, as well as the technical artefacts present in patient tumors coming from different sources (CMI, MET500 and TCGA), producing an integrated cancer expression space with clear clusters composed of mixture of cell lines, xenografts and patient tumor samples. Fig. 2c shows that the aligned expression profiles largely cluster together by disease type, even though MOBER does the alignment in a completely unsupervised manner, without relying on any sample annotations such as disease type. We quantified this by classifying the most similar tumor type for each cell line and xenograft model, based on its nearest neighbors among the TCGA tumor samples. We found that, for 73% of the PTX models the inferred disease type matches the annotated PTX tumor type, while this number goes down to 53% for CCLE models (see Methods).

A key advantage of MOBER is that it does not assume that any two datasets are necessarily similar to each other, and it does not rely on clinical annotations of individual tumor samples, independently of whether they come from patient biopsies or pre-clinical models. As a result, the MOBER-aligned expression data can be used to identify which pre-clinical models have the greatest transcriptional fidelity to clinical tumors, and which models are transcriptionally unrepresentative of their respective clinical tumors. Although a high proportion of pre-clinical models clusters with tumors of the same cancer type, not all cell lines and xenograft models align well with patient tumor samples. Fig. 2d,e show that a significant proportion of cancer models (both PTXs and CCLEs), derived from skin, colorectal and breast cancer are faithful representatives of patient tumors, while many models derived from liver, esophagus and bile duct tend to align with other cancer types. This observation is in agreement with Celligner results on cancer cell lines^19^. Interestingly, PTX models derived from brain tumors align very well with brain cancer patient biopsies, however, brain cell line models cluster closely to soft tissue patient tumors, but not to clinical brain tumors. This is in line with previous studies that show that *in vitro* media conditions cause genomic alterations in brain cell lines that were not present in the original tumors, thus altering their phenotypes^30,31^.

Interestingly, metastatic tumors from the MET500 dataset tend to cluster together with their corresponding primary tumors from TCGA, even though the tissue of biopsy of MET500 tumors is different from the primary site (Suppl. Fig. 2). In 63% of MET500 samples, the inferred disease type matches the annotated primary tumor type. We note that for 88 samples from the MET500 dataset, the primary site annotation is missing (such samples are shown in black squares in Suppl. Fig. 2). However, the majority of these samples align nicely within TCGA clusters, indicating that the unsupervised pan cancer alignment with MOBER can be used to infer the primary site for tumors of unknown origin when the transcriptomics profiles are available.

Another key feature of MOBER is that it allows for populations that are only present in one dataset to be aligned correctly in an unsupervised manner. For example, the CMI dataset contains only transcriptional profiles of patient biopsies with metastatic breast cancer. MOBER integrated this dataset very nicely with the other breast patient tumors from TCGA and MET500, as well as breast cancer cell lines and xenograft models (Fig. 2b).

### MOBER alignment preserves biological subtype relationships

We next sought to determine whether the MOBER alignment keeps known biological differences between more granular cancer subtypes. Focusing here on the breast cancer samples, we determined subtype annotations with the PAM50 method (see Methods). Fig. 3a shows that breast cancer patient biopsies, breast cell lines and xenograft models primarily cluster together by breast cancer subtype (LumA, LumB, Normal, HER2+ or Basal), with only a few PTX Basal subtype models clustering elsewhere (see also Suppl. Fig. 3).

**Fig. 3.**
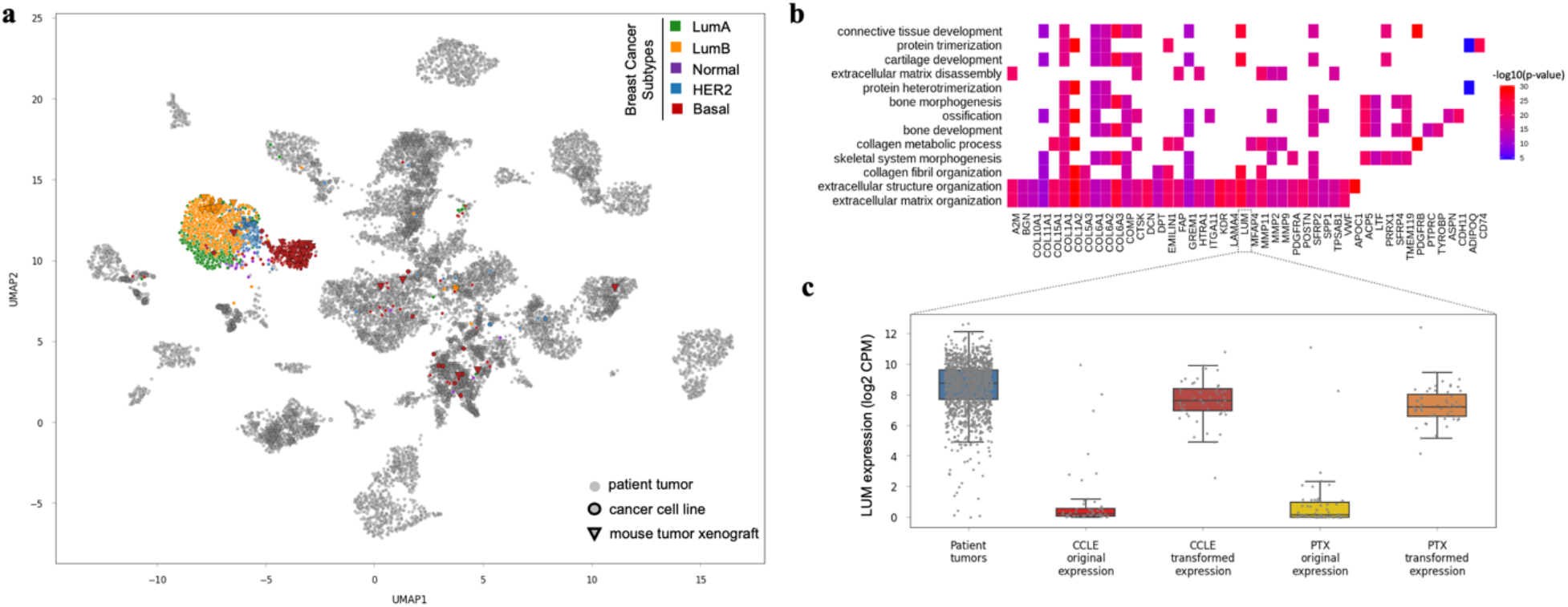
Alignment of breast cancer subtypes. a) UMAP 2D projection of the MOBER-alignment highlighting breast tumor samples: LumA (green), LumB (orange), Normal (purple), HER2 (blue) and Basal (red). All other non-breast tumor samples are in gray. b) Genes that were most significantly upregulated *in silico* after the alignment of breast cancer cell lines to breast cancer patient biopsies (x-axis), along with top enriched biological pathways involving the 100 most upregulated genes (y-axis). c) Expression values (log2 CPMs) of a selected gene, *Lumican (LUM)*, in breast cancer patient tumors (blue), breast cancer cell lines before the alignment (bright red), breast cancer cell lines after the MOBER alignment (dark red), breast xenograft models before the alignment (yellow) and breast xenograft models after the MOBER alignment (orange).

MOBER is fully interpretable by design, which allows us to study the changes in the transcriptomics profiles of pre-clinical models after their *in silico* transformation to clinical tumors. In this respect, we examined the genes that were significantly changed when breast cancer cell lines were aligned to patient biopsies. Fig. 3b shows the genes that were most significantly upregulated after the alignment, along with enriched biological pathways involving these genes. Almost all top enriched biological pathways (e.g. *extracellular matrix organization, collagen fibril organization, connective tissue development* etc) are related to the high presence of stromal tumor microenvironment in breast cancer patient biopsies, that is missing in cell lines (see also Suppl. Fig. 4a). Fig. 3c illustrates the changes in the expression values of a selected gene, *Lumican (LUM)*, in pre-clinical models before and after their *in silico* transformation. *Lumican* is involved in *extracellular matrix organization* and *connective tissue development processes* (among others), and is highly expressed in stromal components in breast cancer patient biopsies, while it has a low expression in pre-clinical models as they are missing the human stromal component. After transforming the breast pre-clinical models to resemble breast patient biopsies, we see that the *LUM* expression was increased in the transformed data. Similarly, when transforming blood cancer cell lines to patient biopsies, the most significantly upregulated genes after the alignment, are related to the presence of human immune components in blood cancer patient biopsies, that are missing in cell lines (Suppl. Fig. 4 b & c). Together, these results highlight that during the projection of pre-clinical models to clinical biopsies, MOBER does not correct genes at random, but it upregulates *in silico* the genes that are expressed in the human tumor microenvironment, which is present in clinical biopsies, but missing in pre-clinical models, while still preserving the key biological information on tumor subtypes at a very granular level.

### Information transfer between cell line and patient tumor datasets

Next we demonstrate how the MOBER-transformed gene expression profiles of pre-clinical models into clinical tumors can be utilized in other studies where we seek to translate pre-clinical biomarkers to patients. The Broad Institute recently published a large metastasis map dataset (MetMap)^15^, determining the metastatic potential of ∼500 human cancer cell lines, thus enabling the metastatic patterns of cell lines to be associated with their genomic features. Here, we sought to identify transcriptomics features that are associated with high or low metastatic potential in human cancer cell lines. We built Machine Learning (ML) models that take as an input gene expression profiles of cancer cell lines and try to predict their average metastatic potential towards 5 different organs, as provided by the MetMap dataset (Fig. 4a, see Methods). Next, we used the models trained on cell line expression data to predict the metastatic potential scores of patient tumors from TCGA. We examined whether there is as difference in the survival between patients for which we predict high metastatic potential vs low metastatic potential. In addition, we determined if there is any correlation between the predicted metastatic scores and the clinical stage of patient tumors. Using the ML models that are trained on original gene expression profiles of CCLE cell lines we see that there is a significant difference in the survival of TCGA patient tumors for which we predict very high metastatic potential (top 25%) vs low metastatic potential (bottom 25%) (p-value = 1.1e-16) (Fig 4b). However, there is no association with clinical stage of TCGA patient tumors (Fig 4c). Then we built new ML models that can predict the metastatic potential scores, but this time we trained them on gene expression profiles of cell lines that are transformed with MOBER to resemble TCGA tumors. Applying these ML models on TCGA patient tumors, we achieve more significant survival stratification of TCGA patient tumors for which we predict very high metastatic potential (top 25%) vs low metastatic potential (bottom 25%) (p-value = 6.2e-29) (Fig. 4d). In addition, we note that such models predict higher metastatic potential of the late-stage tumors, compared to early-stage tumors (Fig. 4e). The ML models that we used here might be too simple to faithfully infer the metastatic potential of tumors, however our results demonstrate the utility of using the MOBER-transformed gene expression profiles in finding biomarkers that are better translatable to patients.

**Fig. 4.**
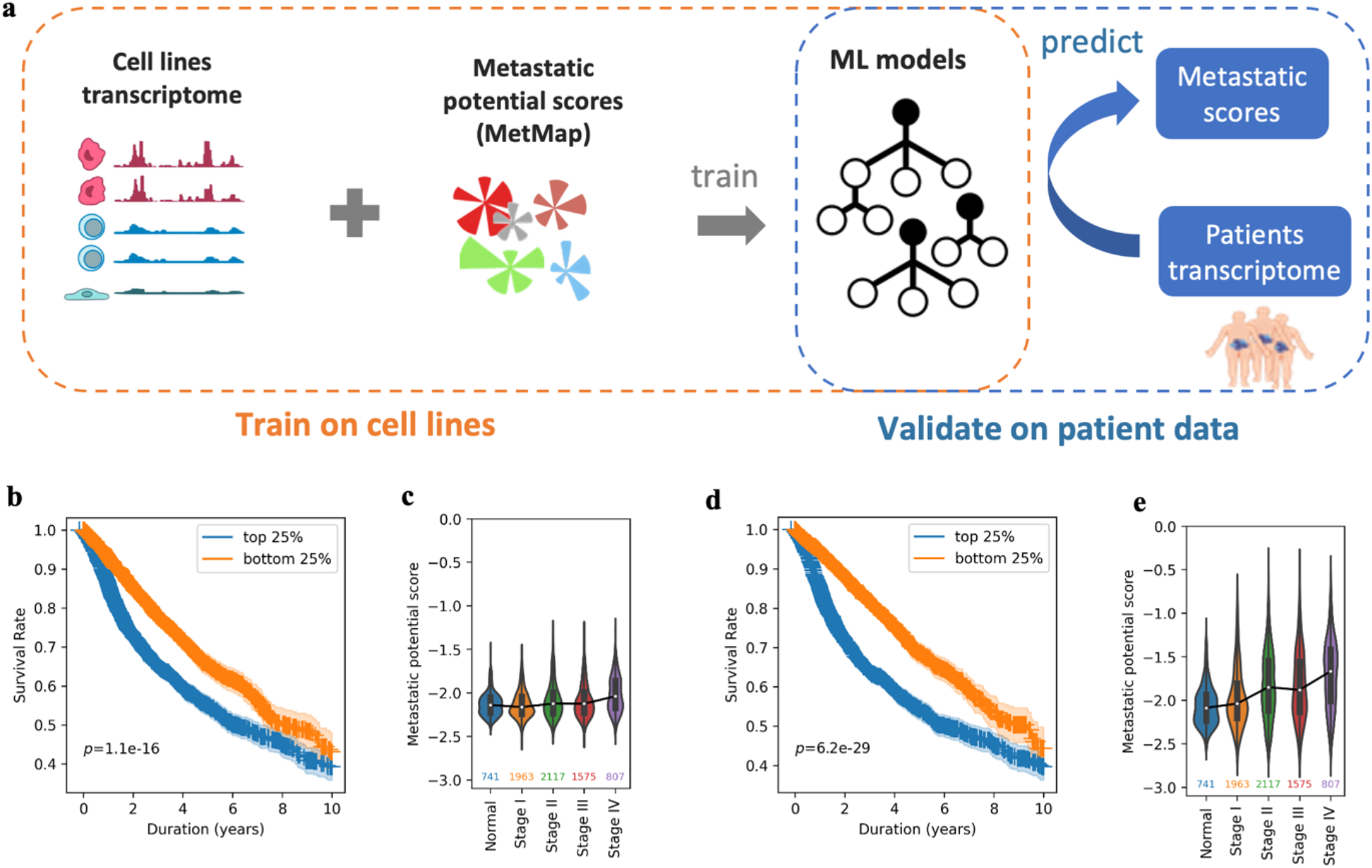
Associating biomarkers of high/low metastatic potential in human cancer cell lines from MetMap and translating them to patients. a) Using gene expression profiles of cancer cell lines and corresponding metastatic potential scores from the MetMap data, we built machine learning (ML) models to find features associated with high/low metastatic potential. Then we translated these models to patients, trying to predict metastatic potential of TCGA patient tumors using gene expression profiles from patient biopsies. b) and c) show that when ML models are trained on original cell line expression profiles from the CCLE, these models transfer poorly to patients. b) Difference in survival of TCGA patient tumors for which we predict very high metastatic potential (top 25%, blue) vs low metastatic potential (bottom 25%, orange) with ML models trained on original cell line expression profiles. P-values are derived from the log-rank test, shaded areas indicate 90% of confidence intervals. c) Predicted metastatic potential of TCGA tumors for different clinical stages. d) and e) show the same as b) and c), but with ML models trained on MOBER-transformed cell line expression profiles to resemble TCGA patients. These models translate better to patient tumors.

## Discussion

Pre-clinical cancer research critically relies on *in vitro* and *in vivo* tumor models, such as cell lines and patient-derived tumor xenografts. However, due to the differences in the growing conditions and the absence of the human stromal microenvironment, these models are often not predictive of the drug response in clinical tumors and do not follow the same pathways of drug resistance^10,32^. Therefore, the identification of the best models for a given cancer type, without relying on annotated disease labels is critically important. To address this, we developed MOBER, a deep learning based method that performs biologically relevant integration of transcriptional profiles from various pre-clinical models and clinical tumors. MOBER can be used to guide the selection of cancer cell lines and patient-derived xenografts and identify models that more closely resemble clinical tumors. We integrated gene expression data from 932 cancer cell lines, 434 patient-derived tumor xenografts and 11’159 clinical tumors simultaneously and demonstrate that MOBER can conserve the inherent biological signals, while removing confounder information.

We identified pronounced differences across cancer models in how well they recapitulate the transcriptional profiles of their corresponding tumors in patients. Certain cancer types, such as the ones in skin, breast, colorectal, kidney and uterus exhibit greater transcriptional similarity between models and patients, while models of cancers of the bile duct, liver and esophagus have transcriptional profiles that are unlike their patient tumors. There are also striking differences between CCLE and PTX as cancer model systems. While brain and soft tissue CCLEs appear to have diverged from their corresponding patient tumors, brain and soft tissue PTX models have high transcriptional fidelity. Notably, among the PTX models, urinary tract xenografts attain greatest transcriptional similarity to corresponding patient tumors, however, many of the urinary tract CCLEs have drifted away from their labeled tumors. As pointed in previous studies^19,33^, many CCLEs are not classified as their annotated labels. We report that PTX models, on average, show greater transcriptional similarity to patient tumors, as compared to their CCLE counterparts. This could suggest that the lack of immune component is not the main CCLE confounder, but CCLEs likely undergo transcriptional divergence due to the culture condition, high number of passages and genetic instability^34,35^. In addition, cancer models (both PTXs and CCLEs) could be misannotated due to inaccurate assignment based on unclear anatomical features or mismatch during sampling^33,36^. Our pan cancer global alignment with MOBER could be used to identify cancers that are underserved by adequate pre-clinical models, and to determine the primary site label for models and clinical tumors where such annotation is missing.

MOBER is interpretable by design, therefore allowing drug hunters to better understand the underlying biological differences between models and patients that are responsible for the observed lack of clinical translatability. The observed differences vary across disease types, emphasizing the importance of using unsupervised nonlinear approach that enables identification of disease-type-specific variations.

As a batch effect removal method, MOBER offers several advantages compared to previously published methods^19,21,37^: i) supports integration of multiple datasets simultaneously; ii) enables transformation of one dataset into another; iii) does not make any assumption on the datasets composition. We demonstrate that MOBER can remove batch effects between any gene expression datasets, even when the cell population representation across datasets is different.

In order to facilitate the use of MOBER, we made the source code available at https://github.com/Novartis/MOBER, and developed an interactive web app available at https://mober.herokuapp.com to allow users to explore the MOBER aligned expression profiles coming from cancer models and clinical tumors. In addition, the web app enables the identification of pre-clinical models that best represent the transcriptional features of a tumor type, or even a particular tumor subtype of interest. Future version of MOBER that integrates genetic and epigenetic features of models and patient tumors could potentially enable even more detailed analysis between models and patients.

## Methods

### The MOBER method and model training details

Each gene expression profile of a sample *i* is a vector *x*_*i*_ with length equal to the total number of genes. The input gene expression data *x* is run through a variational autoencoder (VAE) that uses variational inference to reconstruct the original data in a conditional manner. The encoder estimates the probability density function of the input expression data *Q*(*z*|*x*). Then, a latent vector *z* is sampled from *Q*(*z*|*x*). The decoder decodes *z* into an output, learning the parameters of the distribution *P*(*x*|*z*). The loss function is then given by:

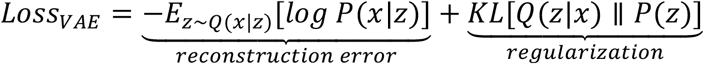

where *x* and *z* indicate the gene expression data and latent space respectively, *Q* and *P* are the estimated probability distributions, *E* denotes an expectation value and *KL* is the *Kullback-Leibler*^38^ divergence. The first term in the loss function is the reconstruction error (i.e. expected negative log-likelihood of the data sample), and the second term is the *Kullback-Leibler* divergence between the encoder’s distribution *Q*(*z*|*x*) and *P*(*z*). In addition to the input expression data, we provide to the decoder an information about the origin of the input sample (in our case CCLE, PTX, TCGA, MET500 or CMI) transformed into a one-hot encoding vector *s*, that consists of 0s in all cells and a single 1 in a cell used to uniquely identify the input source. This allows for reconstruction of the latent vector *z* by the decoder in a conditional manner, and it enables projection of one dataset into another.

In addition to training the VAE, we simultaneously train an adversarial neural network (aNN) that acts as a source discriminator. It takes as an input an embedding vector *z* sampled from the latent space, and tries to predict the source label of the input data (*s*). This is a multi-class fully connected neural network with negative log-likelihood loss function as given by:

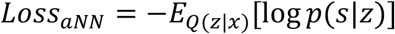

The joint loss is computed as:

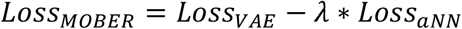

where λ is a coefficient that determines the weight that the model gives to the adversarial loss.

In our study, the encoder, decoder, and aNN are designed as fully connected neural networks each with 3 layers. Each layer of the encoder (and decoder respectively) consisted of 256, 128 and 64 nodes each. We used *Scaled Exponential Linear Units* (SELU)^39^ activation function between two hidden layers, except the last decoder layer, where we applied Rectified Linear activation Unit (ReLU)^40^. The last aNN layer had 5 hidden nodes corresponding to the number of data source classes and softmax activation.

We implemented MOBER using *PyTorch*^41^. We set the minibatch size to 1600, and trained it with Adam optimizer^42^ using a learning rate of 1e-3. The weight for the KL loss of the VAE was set to 1e-6, the weight for the source adversary loss was set to 1e-2. The best hyper-parameter set from numerous possibilities was chosen from a grid search that minimized the joint loss and maximized the clustering performance of models to patients.

### Datasets

The RNAseq gene expression counts for TCGA samples were downloaded from the TCGA portal (https://tcga-data.nci.nih.gov/tcga)^14^, the CMI gene expression counts from the Genomic Data Commons (GDC) portal^43^, and CCLE^2^, PTX^9^ and MET500^29^ data were obtained from the corresponding publications. All gene expression data were TMM-normalised using EdgeR^44^ and then transformed to log2 counts-per-million using the *edgeR* function ‘cpm’, with a pseudocount of 1 added. Gene expression data were subset to 17’167 protein-coding genes that were present in all datasets. The MetMap^15^ dataset was downloaded from the DepMap portal (https://depmap.org/metmap). We excluded the indications that have less than 5 samples.

### Alignment evaluation

We projected each sample from the CCLE, PTX, CMI and MET500 datasets to TCGA, by changing the one-hot encoded source information and setting it to TCGA. Then we decoded the expression data with MOBER.

To evaluate the alignment of pre-clinical samples to TCGA samples, for each CCLE and PTX sample we identified the 25 TCGA nearest neighbors in 70-dimensional PCA space. Each pre-clinical sample was classified as a tumor type by identifying the most frequently occurring tumor type within these 25 nearest neighbors.

The identification of breast cancer molecular subtypes was done using the PAM50 classifier as implemented in the *geneFu* R package^45^. The identification of differentially expressed genes comparing the transcriptional profiles before and after their transformation to patient tumors was done with *Seurat* v3.6^46^ using the t-test method. Pathway enrichment analysis was done for the top 100 most differentially expressed genes ordered by their fold change and with an adjusted p-value < 0.01 using the *clusterProfiler*^47^ package.

### Analyses of metastatic potential using MetMap data

We trained random forest models to predict the metastatic potential scores using gene expression values as input, using the *scikit-learn*^48^ Python package (0.23.2). The hyperparameters were optimized with grid search strategy using 3-fold cross validations. Then the final model was trained using the optimized hyperparameters. The hyperparameter optimized in the model is “max_features”. 1000 trees were used, and all other hyperparameters were set as default.

Two different models were trained, using either the original transcriptome or the projected transcriptomes of CCLEs to TCGA patients to predict mean metastatic potential scores of cell lines across 5 organs (*metp500*.*all5*). Then with each trained model, we predicted the metastatic potential scores for patient tumors from TCGA using the original transcriptome as input. For the survival analysis, the top 25% of samples with highest predicted scores and bottom 25% with the lowest predicted scores were compared. The *Kaplan-Meier* survival analysis^49^ was done with the *lifelines*^50^ Python package v0.25.4.

## Code availability

The source code of the MOBER method is available at https://github.com/Novartis/MOBER. An interactive web app to explore the MOBER alignment in details is available at https://mober.herokuapp.com.

## Acknowledgements

G. L. would like to acknowledge the support from Novartis Innovation Postdoctoral Fellowship program.

## Supplementary Figures

**Suppl. Fig. 1.**
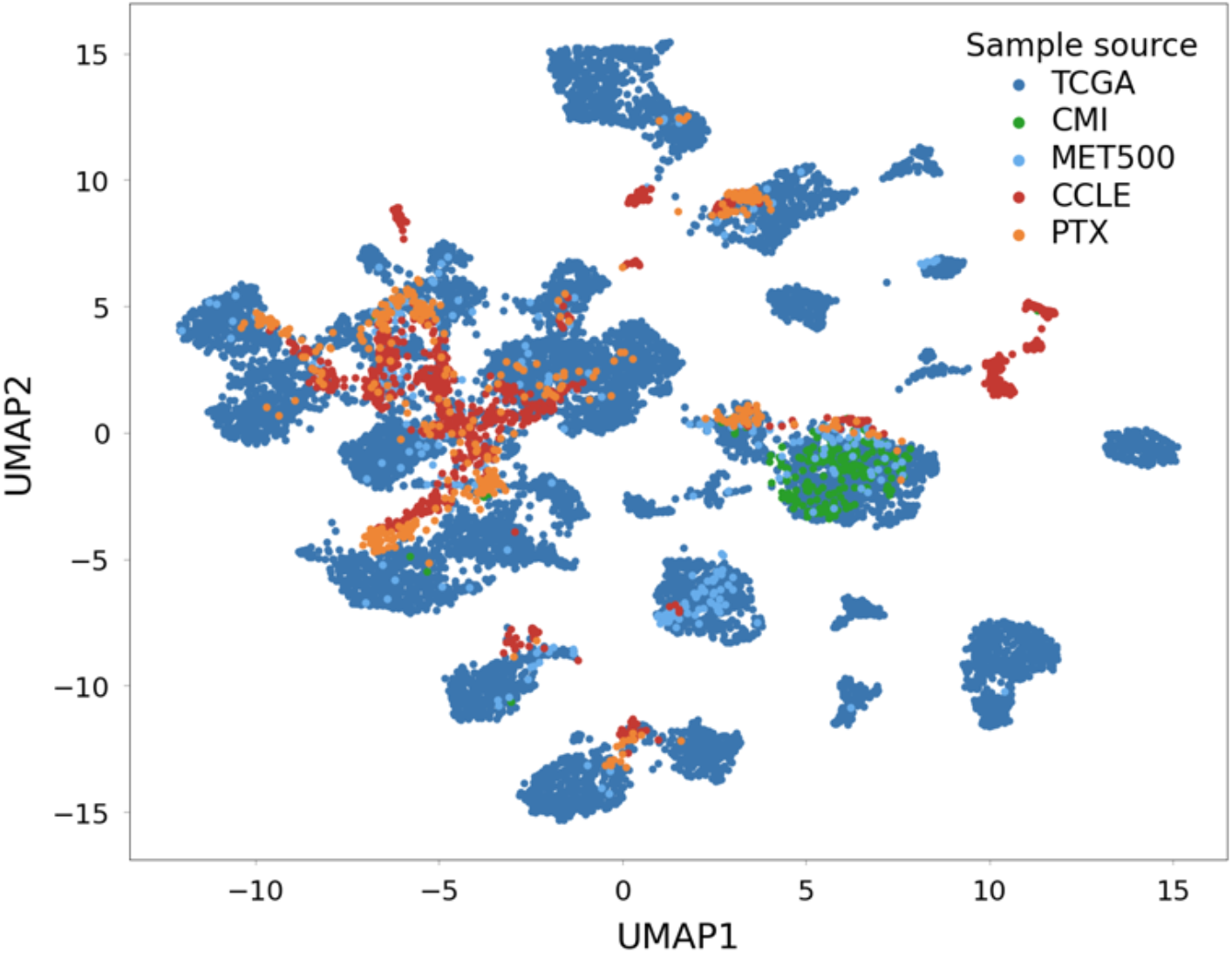
Batch effect correction with ComBat.

**Suppl. Fig. 2.**
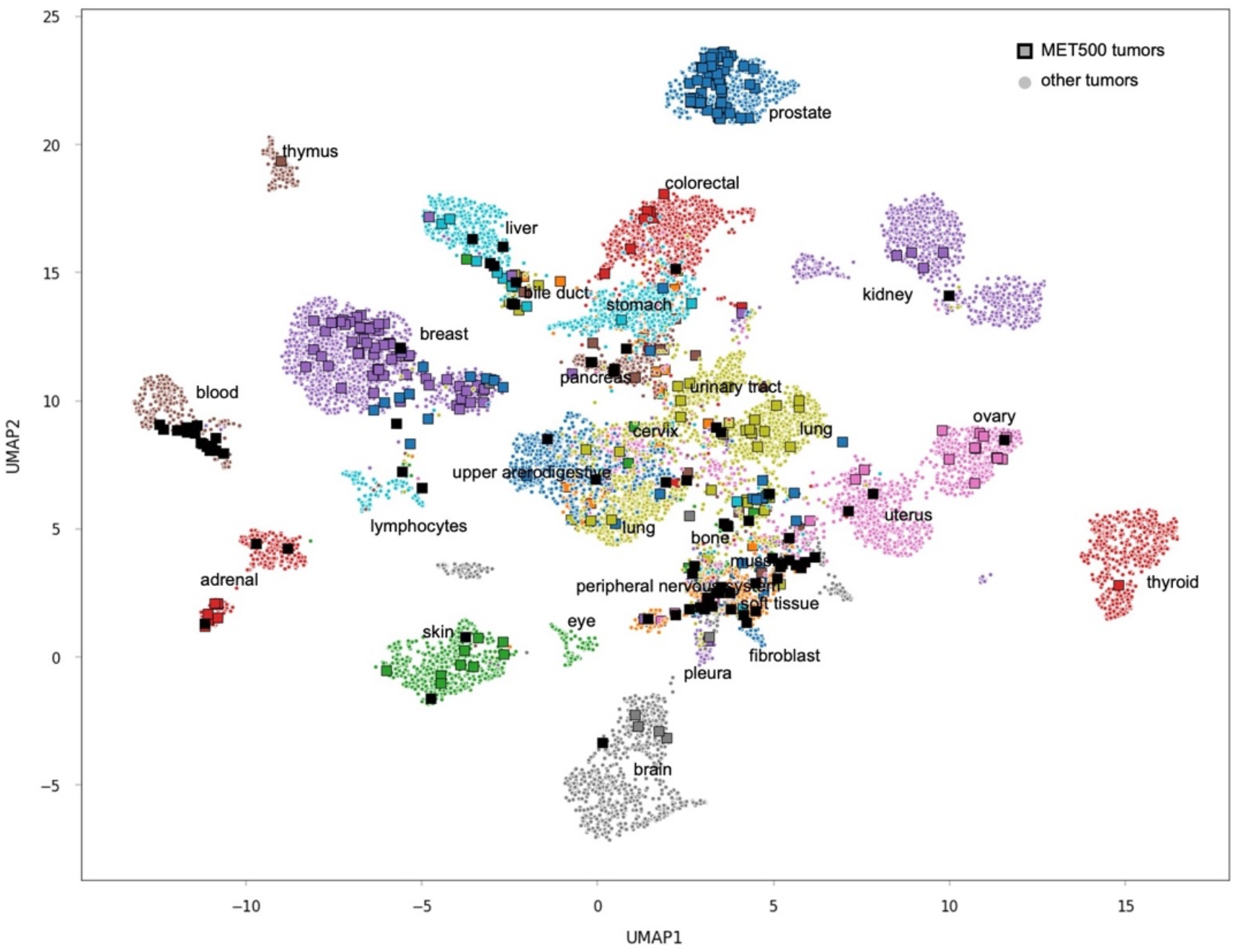
Global pan cancer alignment with MOBER, highlighting MET500 metastatic samples. MET500 tumors are shown in squares with color corresponding to the primary site. The samples with unknown primary origin are shown in black. All other tumors (CCLE, PTX, CMI and TCGA) are shown with circles and are colored based on their primary site annotation.

**Suppl. Fig. 3.**
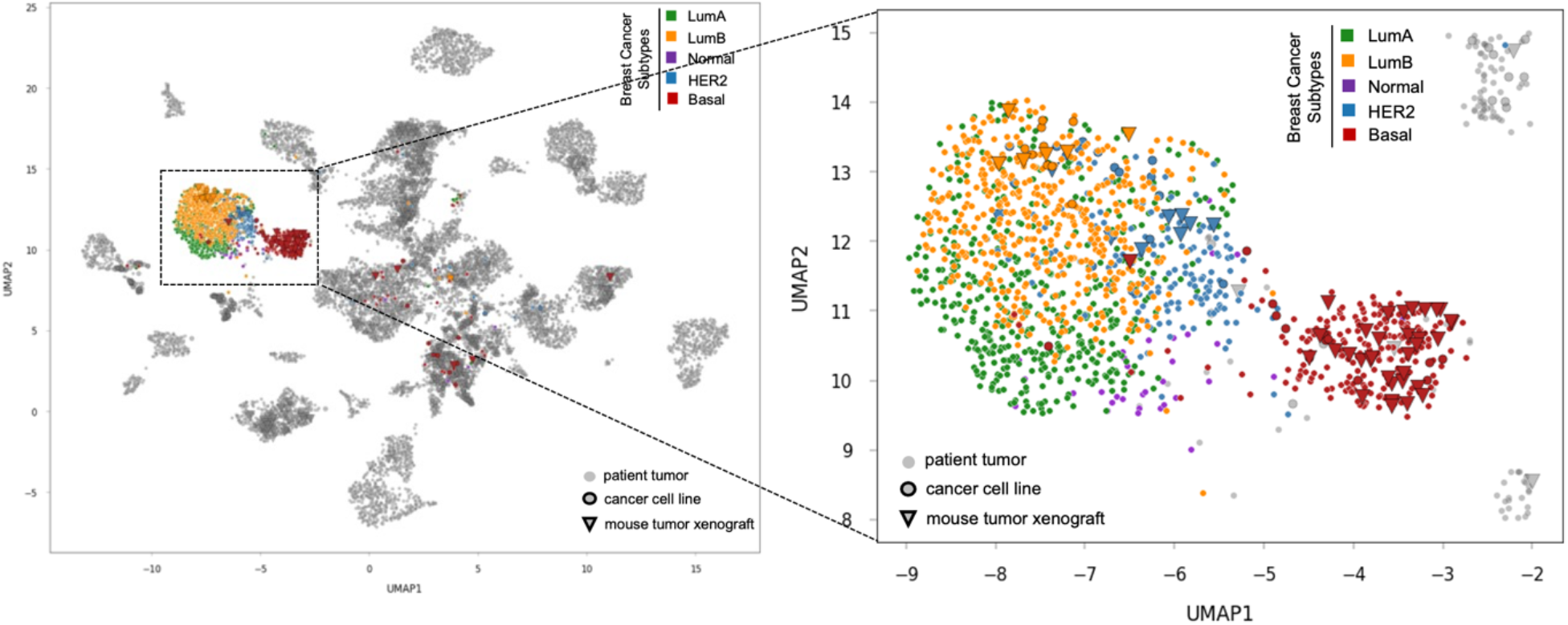
Alignment of breast cancer subtypes. UMAP 2D projection of the MOBER-alignment highlighting breast tumor samples: LumA (green), LumB (orange), Normal (purple), HER2 (blue) and Basal (red). All other non-breast tumor samples are in gray.

**Suppl. Fig. 4.**
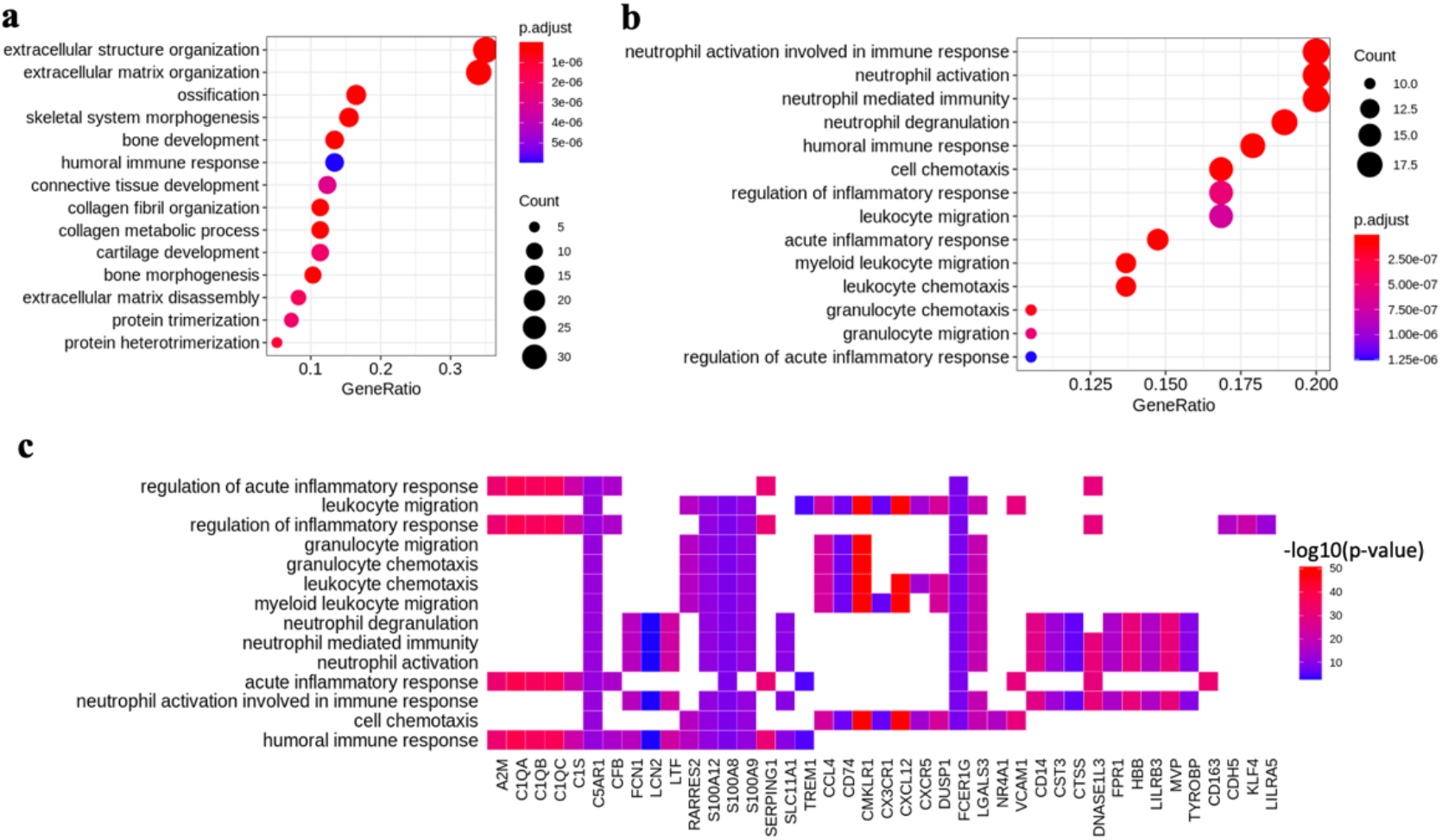
Pathway enrichment analysis of the top 100 upregulated genes when transforming CCLEs to TCGA tumors for a) breast primary tumors; b) blood tumors. c) Genes that are most significantly upregulated *in silico* after the alignment of blood CCLEs to blood TCGA tumors (x-axis), along with top enriched biological pathways involving the 100 most upregulated genes (y-axis).

